# Discriminating tactile speed in absence of raised texture elements: Role of deformation and vibratory cues

**DOI:** 10.1101/599522

**Authors:** Alessandro Moscatelli, Colleen P. Ryan, Simone Ciotti, Lucia Cosentino, Marc O. Ernst, Francesco Lacquaniti

## Abstract

Motion encoding in touch relies on multiple cues, such as displacements of traceable texture elements, friction-induced vibrations, and gross fingertip deformations by shear force. We evaluated the role of deformation and vibration cues in tactile speed discrimination. To this end, we tested the discrimination of speed of a moving smooth glass plate, and compared the precision of the responses when the same task was performed with a plate having a fine texture. Participants performed the task with and without masking vibrations. Speed discrimination was nearly as precise among the two surface types, as assessed by the steep slope of the psychometric function. Consistent with our previous work, high-frequency vibrations impaired the ability of the participants in discriminating surface speed. Results of the current study showed that it is possible to discriminate motion speed even in absence of a raised texture.

**Highlights:**

- On a smooth surface, humans are able to discriminate the speed of a moving surface by frictional motion cues
- The precision of speed discrimination is nearly the same with smooth and fine-textured surface types
- High frequency vibrations impair the ability to discriminate speed of moving surfaces

## Introduction

It has been argued that motion is as important to touch as light is to vision [1]. Tactile motion plays a central role in perceptual and motor tasks. It provides feedback for the manipulation of handheld objects [2], and for guiding hand reaching towards a target goal while touching a surface [3, 4]. During grasping tasks, our sensorimotor system rapidly adjusts the grip force based on the small slips between the object and the skin, revealed as vibrations [5].

Different cues contribute to the perception of motion in touch, including the spatiotemporal pattern of skin indentation, as for example the one generated by the tip of a pencil moving across the skin, high-frequency vibrations caused by frictional slips, and the gross deformation of the skin and subcutaneous tissues, such as the stretch generated by a shear force applied to the fingertip [6, 7, 8, 9, 10]. The role of the first two motion cues, local indentation and vibrations, was investigated in several studies [11, 12, 13, 14, 15, 16, 17, 18]. Here, we investigated the discrimination of speed of a smooth movable surface (i.e., a glass plate for microscope slide). Because of the lack of fine and coarse texture, shear deformation cues may play an important role for this type of stimuli. Before introducing our study, we will summarize recent findings about different cues in tactile motion perception.

The indentation produced by traceable surface elements, like raised ridges of the tip of a pen, provide a salient motion cue in touch [6, 11, 13]. For instance, the perceived speed of a ridged surface depends on the distance between its ridges, such as the closer the texture elements, the faster is the perceived speed [12, 19]. In accordance with behavioral studies, a model based on the spatiotemporal pattern of skin deformation, as the one produced by a coarse texture, accurately reproduced tactile afferent signals [20]. In addition, the somatosensory system uses heuristic cues arising from frictional skin vibrations that are generated by sliding across or tapping against the surface of an object [21, 14, 16]. Frictional vibrations play an important role in motion detection, speed discrimination, and roughness discrimination [6, 22, 17, 18]. Their importance as a motion cue has been demonstrated mostly during slip motion on fine texture surfaces, lacking traceable elements, while their role is likely to be less relevant on surface with coarse textures [6, 17]. A third type of motion cues are referred to as gross deformation cues, such as the tangential stretch of the skin caused by shear force and the transition from stick to slip between the skin and a surface [6, 10, 23]. A smooth glass plate, lacking fine and gross texture elements, has been used to investigate deformation cues [6, 10]. For light touch, i.e., for a normal force equal to 0.2 N, the slip of a smooth glass plate is indistinguishable from simple skin stretch [6]. Instead, it is possible to detect slip and partial slip of a glass surface at higher values of normal force, ranging from 2 to 5 N [10]. These behavioral results complement electrophysiological recordings of the afferent fibers. A moving glass plate elicits only a weak response in fast adapting fibers during light touch [6], whereas, for a normal force up to 4 N, nearly all afferents respond to tangential strain, and their response is broadly tuned to a preferred direction of force [24].

Here, we aimed to further investigate the role of deformation and vibration cues for motion encoding in touch. To this end, we tested the discrimination of tactile speed on a smooth glass plate. From the precision of the response, it is possible to estimate to which extent participants were able to discriminate surface speed in absence of raised texture elements. For comparison with previous studies, the task was replicated using a surface with a homogeneous texture similar to the one used by [17, 25]. In both surface conditions, high-frequency vibrations were delivered synchronously with surface motion in half of the trials. This was to investigate the possible interplay between deformation and vibration cues. If these two cues were processed in separate channels, and speed discrimination on a smooth plate depended on deformation cues only, masking vibrations should affect the precision of the response on the textured but not on the smooth plate. Otherwise, this would speak in favor of an interplay between vibration and deformation cues, which may occur at a mechanical or at a neural level. The role of contact force in this task was investigated systematically in a second experiment.

## 1. Experiment 1: Speed discrimination on smooth and textured surfaces

In the first experiment, we investigated the speed discrimination on a movable smooth surface (microscope glass plate) and compared these results against the speed discrimination on a fine textured surface (3D-printed Polylactic Acid plate). The effect of masking vibrations in the two surface conditions was also assessed.

### Methods

#### Participants

Fifteen naïve, right-handed participants took part in the experiment (10/15 females, age ranging from 20 to 36 years; 13 naïve participants plus author AM and CPR). The testing procedures were approved by the ethics committee of the Santa Lucia Foundation, in accordance with the guidelines of the Declaration of Helsinki for research involving human subjects. Informed written consent was obtained from all participants involved in the study.

#### Apparatus

A motion device displaced the contact surface up and down at the required speed (Figure 1A-B). The motion device consisted of two rollers with a rigid plane in between, rotating a plastic belt (270 mm length × 30 mm width). The two rollers were vertically aligned and the downward roller was connected with a metal shaft to a Faulhaber motion control system. This included a DC-Micromotor (Faulhaber 3242G024CR combined with 7:1 Gearheads 32A), a high-resolution encoder (Faulhaber IE3-512), and a position and speed controller (Faulhaber MCDC3006SRS). A rigid plastic box having a cuboid shape (75 × 20 × 20 mm) was attached to the belt. This way, the rotation of the motor produced a vertical displacement of the box. We assembled two boxes having a contact surface of different materials, glass and plastic. The base of each box was interlocking with a basement firmly attached to the belt, so that they could be easily changed between two experimental blocks. The contact surface was either a smooth glass plate (75 × 25 × 1.5 mm) or a plastic plate (75 × 20 × 1.5 mm). The glass plate was the one used for microscope slides and had no detectable features on its surface. The plastic plate was printed in Polylactic Acid (PLA) using a 0.4 mm diameter nozzle, which produced a fine homogeneous texture on its surface. To detect contact, a force resistive sensor was placed between the box and the contact surface (Interlink 402 force resistive sensor connected to a Arduino UNO microcontroller). Prior to the experiment, the contact sensor was calibrated by means of a set of calibration weights, ranging from 0.98 to 4.9 N. For a more accurate measurement of contact force, the FSR was replaced with a Load Cell in Exp. 2 (Fig. 1B). A vibromotor (Haptuator Mark II by TactileLabs, connected to MAX9744 audio amplifier by Adafruit) was placed in a small niche inside the box to generate masking vibrations. Vibration stimuli were generated by a standard PC audiocard (HDA Intel PCH). Masking vibrations were recorded with an accelerometer attached to the contact plate to measure the amplitude and frequency of the signal (Fig. 2). A custom-made Matlab code controlled the vibration and the motion stimuli. A one degree of freedom finger holder was attached to the frame of the setup, supporting the participant’s finger.

**Figure 1:**
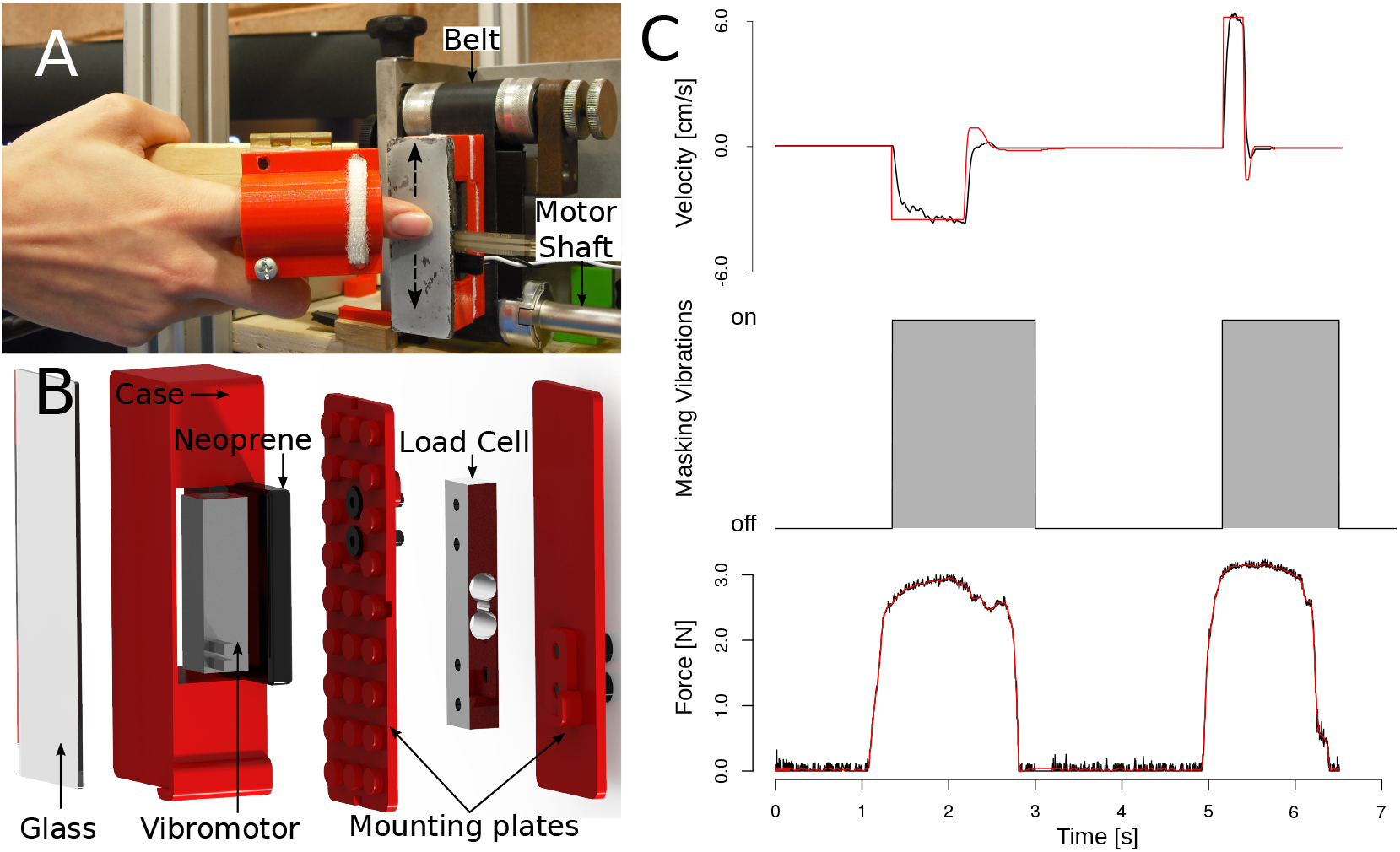
Experimental setup. A) The setup including the actuated box, the vibromotor and the force sensor. Two boxes having a glass or a plastic contact plate were used in two experimental blocks. B) Exploded drawing of the device (modified version used in Exp. 2). C) Motion, vibration and force profile in a single trial. In the upper panel, positive values of velocity correspond to downward motion and negative values to upward motion. Measured and target values are represented in black and in red, respectively. To prevent using motion duration as a cue, path length was randomized across trials. Middle panel: In half of the trials, high-frequency vibrations were presented while the participant contacted the plate (vibration condition). Vibration amplitude was set to zero in the other half of the trials (control condition, not shown in the figure). Vibration onset and offset was synchronized with the initiation and the cessation of contact. The lower panel shows the contact force. Raw and filtered data are represented in red and in black, respectively

**Figure 2:**
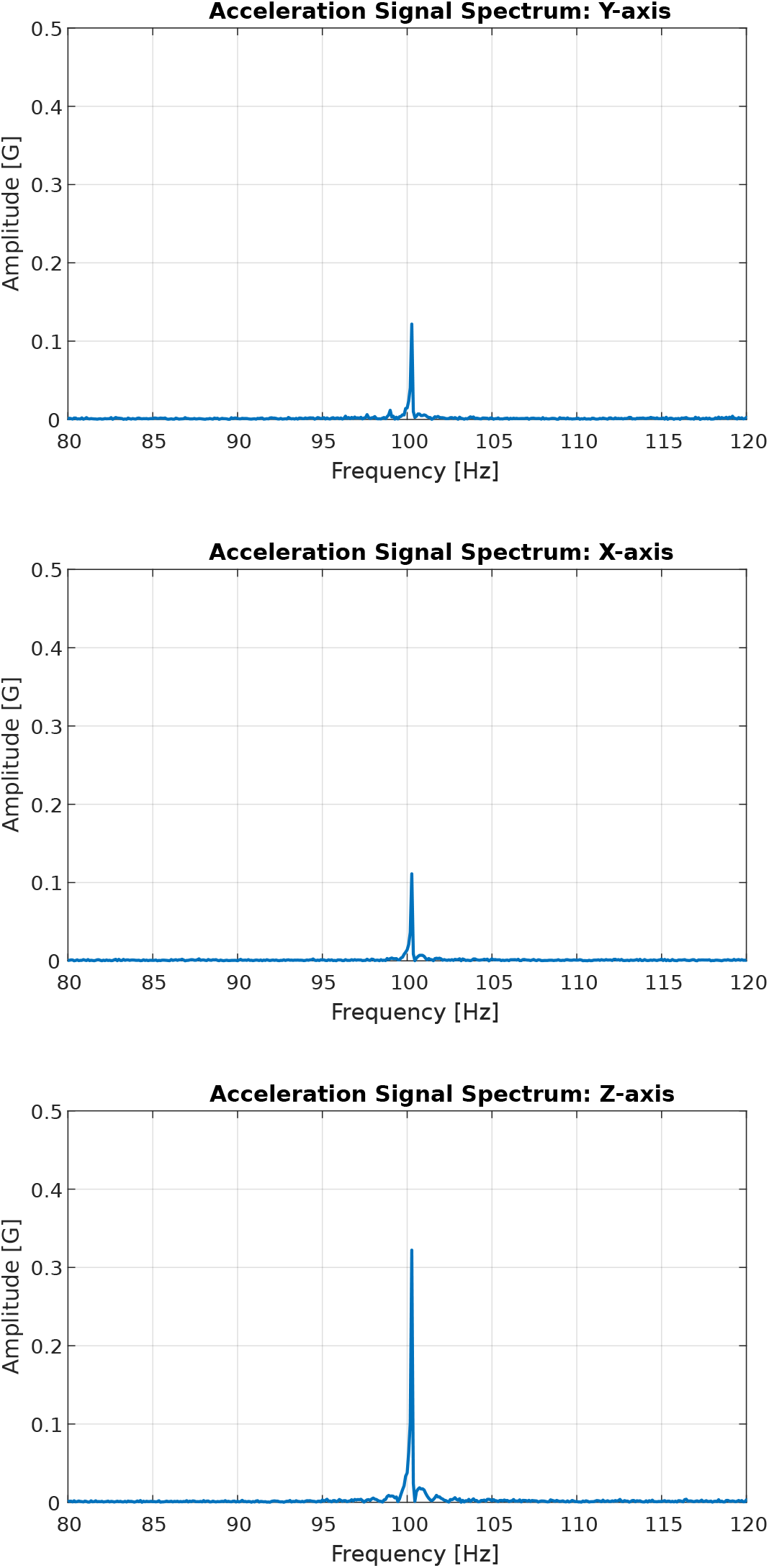
Masking vibrations: x, y, and z acceleration component. Vibrations were recorded with an accelerometer (ADXL337 connected with an Arduino UNO microcontroller, sampling rate 360 Hz) attached to the contact plate, and processed with Fast Fourier Transform.

#### Stimuli and Procedure

Participants sat on an office chair in front of the apparatus, resting their right index finger on the finger holder. A black curtain hid the device from the participants’ sight. Throughout each experimental session, participants wore earplugs and headphones playing pink noise in order to mask external sounds. Participants performed a forced-choice, speed discrimination task. The advantage of using a discrimination task is that, differently from a yes/no detection task, the response is less bias-prone, i.e., it does not depend on the decision criterion. Each experimental session consisted of 240 trials divided into two blocks. In each block, participants performed the experiment either with the smooth glass plate or with the textured plastic plate. The order of the two blocks was counterbalanced across participants.

The experimental procedure was based on the method of constant stimuli. The sequence of the stimuli in a trial is illustrated in Fig. 3 and Fig. ??. Each trial included a reference and a comparison stimulus. The order of the two was counterbalanced across trials. We instructed participants to push on the contact surface with their index finger to start the tactile stimulus. The servomotor and the vibromotor were actuated when the normal force (averaged across the last 20 ms) exceeded the threshold value of 1.5 N. The surface moved either upward or downward, with the motion direction of the second stimulus always being the opposite of first one. The motion speed was equal to 3.43 cm sec^−1^ in the reference stimulus, and it was pseudo-randomly chosen between five values ranging from 0.63 cm sec^−1^ and 6.24 cm sec^−1^ in the comparison. After each trial, participants reported whether the surface moved faster in the first or in the second stimulus interval. To prevent using motion duration as a cue, the path length of stimulus was pseudo-randomly chosen within a range of 10-14 mm. In half of the trials, masking vibrations consisting of a 100 Hz sinusoidal wave were delivered synchronously with the motion stimulus (Fig. 3).

**Figure 3:**
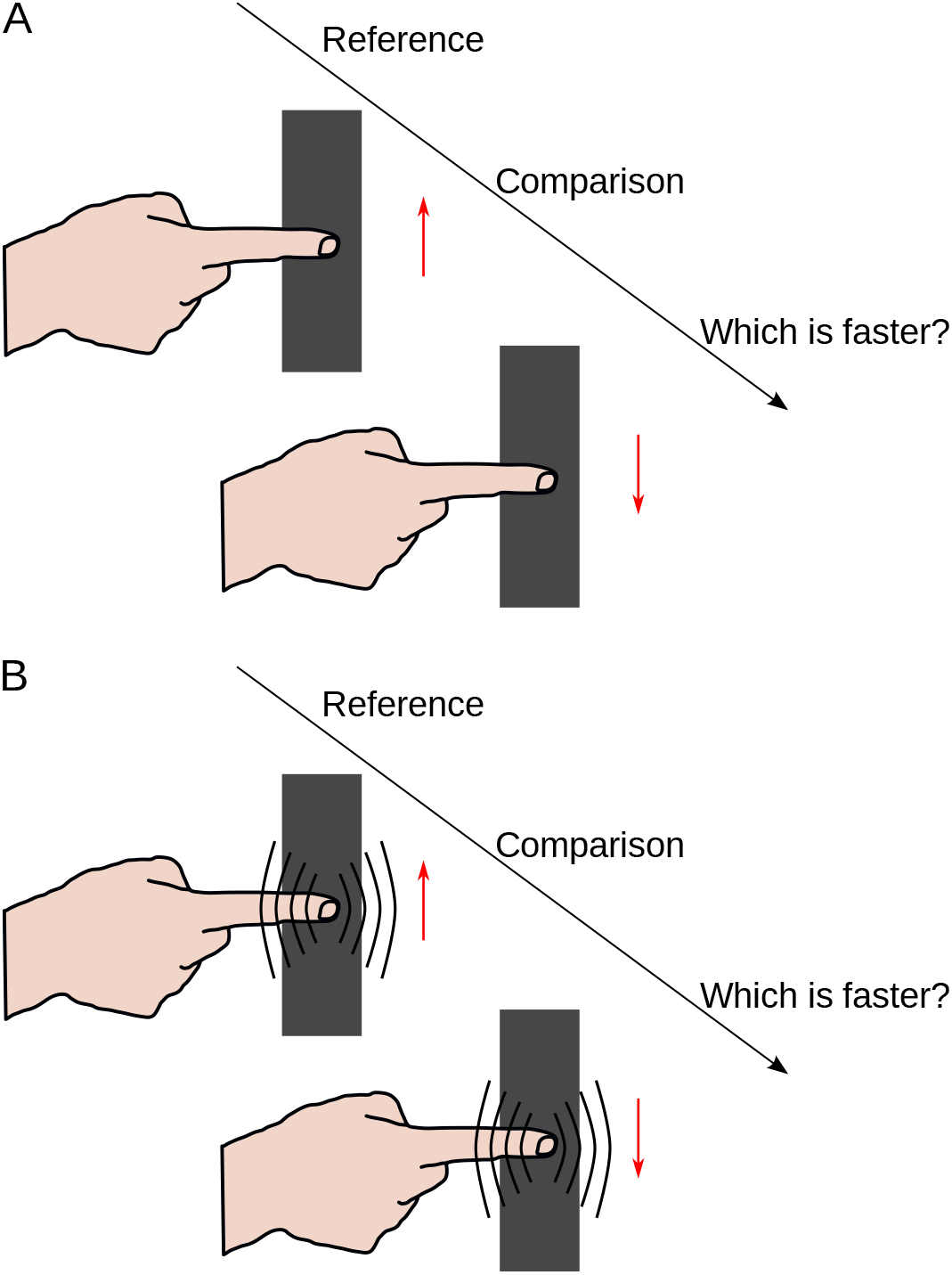
Experimental procedure. A) Control condition B) Masking vibration condition

#### Analysis

We analyzed categorical responses by means of Generalized Linear Mixed Models (GLMM) [26], separately for each surface type, glass and plastic. We included the motion speed (S) and the presence/absence of vibration (V = 0 without vibration and V = 1 otherwise) as fixed-effect predictors.

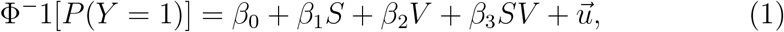

where *P*(*Y* = 1) was the probability of reporting that the comparison was faster that the reference. Fixed-effect parameters were denoted with Greek letter *η*_1_,…,4 and random-effect parameter vector with the lowercase letter 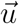. The parameter *β*_1_ (referred to as the model slope) estimates the reliability of the response in the control condition (without vibrations). With vibrations, the model slope was equal to *β*_1_ + *β*_3_. For comparison with previous studies, we computed the Just Noticeable Difference (JND = 0.625/*β*_1_), the Weber Fraction (WF = JND/3.43, which is the value of the reference speed), and the related 95% Confidence Interval (CI) by means of the bootstrap method explained in [26]. Next, we applied the following model to compare the response between the two surface types:

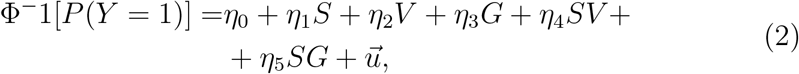

This corresponds to a 2×2 factorial design (2 surface types × 2 vibration conditions). Model (1) and (2) were selected among a pool of nested models based on the Akaike Information Criterion (AIC) and likelihood ratio test (LR test). The R packages *lme4* and *MixedPsy* were used to fit the GLMMs [27, 26].

#### Power Analysis

We performed the power calculation by using the R package *simr* [28]. We assumed a sample size equal to 15 participants, 60 trials each (corresponding to trial repetitions in each experimental condition), and effect size equal to 0.40 (slope of the model, or *β*_2_). With these parameters, statistical power of the GLMM would be above 80%. This sample size is in accordance with the one reported in similar studies in the literature [17, 10, 29].

### Results

The first aim of this study was to test whether participants were able to discriminate the motion speed of the smooth glass surface, with the fine plastic surface serving as control. As illustrated in Figure 4, participants were able to perform the task with either of the two types of surface. Without masking vibrations, the parameter of slope in Eq. (1) was significantly larger than zero with the smooth glass plate (*p* < 0.001; 0.65 ±0.079, *β*_1(*glass*)_ ± Std. Error) and with the homogeneous-textured plastic plate (*p* < 0.001; 0.75±0.10, *β*_1(*plastic*)_ ± Std. Error). This means that, differently from previous studies on the detection of slip motion, participants discriminated the motion speed of the smooth glass plate, well-above chance level.

**Figure 4:**
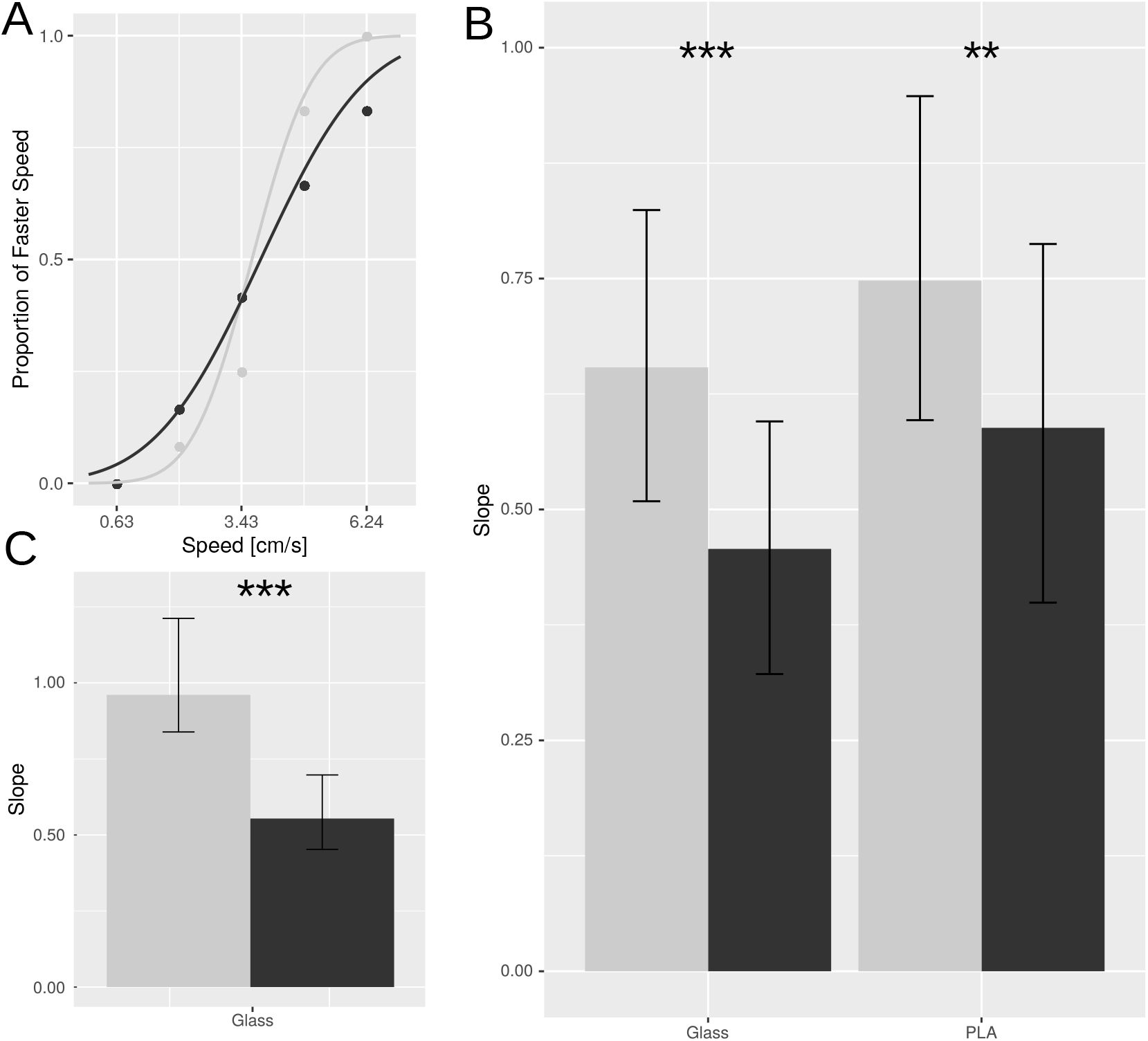
Results. A) Psychometric functions in a representative participant, raw data and GLMM fit (glass surface). Control condition and masking vibrations condition are represented in light and dark grey, respectively. B) The bar plot shows the estimated slopes in the control (light grey) and masking vibration conditions (dark grey) in Exp. 1, separately for the glass and PLA surface (GLMM estimates to 15 participants). The error bar represents the 95% confidence interval estimated with the bootstrap method. C) The bar plot shows the estimated slopes in the control and masking vibration conditions in Exp. 2 (GLMM estimates to 6 participants).

With the glass plate, tactile speed discrimination was impaired by high-frequency vibrations. Accordingly, the slope was significantly smaller with masking vibrations compared to control (*p* < 0.001; −0.20 ±0.048, *β*_3(*glass*)_ ± Std. Error). The same effect of masking vibrations was observed with the plastic surface (*p* = 0.002; −0.16 ±0.05, *β*_3(*plastic*)_ ± Std. Error). Results are illustrated in Figure 4A-B, in a representative participant, and in Figure 4C for the whole population. We evaluated the difference between the two surface conditions by means of the GLMM in Eq. (2). The precision of the response, although higher with the plastic interface, was not significantly different between the two surfaces (*p* = 0.06; 0.057±0.031, *η*_5_ ± Std. Error).^1^

For comparison with previous studies, the values of JND in the four experimental conditions are reported in Table 1, along with the related 95% CI. With the fine plastic surface and in absence of vibrations, the value of Weber Fraction, *WF* = *JND*/*V_ref_*, is equal to 0.26, which is close to the ones reported in previous studies where the skin was stimulated either with a sandblasted surface or with a brush [30, 25, 17].

**Table 1:** Experiment 1: Values of Just Noticeable Difference (JND) in the four experimental conditions. Estimates and 95% CI estimated with the bootstrap methods from the GLMM.

| Estimate | Inferior | Superior | Parameter | Vibration | Surface |
| --- | --- | --- | --- | --- | --- |
| 1.03 | 0.82 | 1.33 | JND | no | Glass |
| 1.48 | 1.13 | 2.10 | JND | yes | Glass |
| 0.90 | 0.71 | 1.13 | JND | no | Plastic |
| 1.15 | 0.86 | 1.69 | JND | yes | Plastic |

## 2. Experiment 2: Evaluating Contact Force

According to previous literature, tactile discrimination of motion may change depending on the level of normal force. For comparison with previous studies, we replicated the task (only on the glass plate) and replaced the force resistive sensor in the apparatus with Load Cell that provides a more accurate measurements of force.

### Participants

Six right-handed participants took part in the experiment (2/6 females, age ranging from 22 to 38 years; 4 naïve participants plus author AM and CPR).

### Apparatus

The apparatus was similar the one in Exp. 1. This time, a load cell (LC, CZL616C Micro Load Cell by Phidgets Inc) combined with LC amplifier (HX711 Load Cell amplifier, sampling frequency: 80 *Hz*) was used to measure contact force (Fig. 1A-B). The specifics of the LC are the following: repeatability error: 390 *mg*; maximum weight capacity: 780 *g*. Force values were read by Arduino UNO by using the HX711 Arduino Library to interface with the amplifier.

### Stimuli and Procedure

The same procedure of Exp. 1 was used. This time, only the smooth glass surface was tested.

### Analysis

As for Exp. 1, the perceptual response of individual participants and population were fitted with GLM and GLMM, respectively. As in our previous study [31], force signals were filtered using a second order, Butterworth low-pass filter (cutoff frequency equal to 10 Hz). An example of raw data and filtered data is illustrated in Fig. 1C. In each trial, we saved the peak of the filtered signal for additional analyses.

### Results

#### Speed Discrimination

Categorical responses were in agreement with the previous experiment. Participants were able to discriminate the velocity of the glass plate; perceptual judgments were less accurate with masking vibrations. In the condition without vibration the slope of the GLMM was 0.96 ±0.10. The estimated value of the slope is significantly different from zero (*p* < 0.0001). The slope in the condition without vibration is significantly steeper from the slope in the condition with vibration, the difference in slope being equal to 0.41 ±0.01 (*p* < 0.0001). This is illustrated in Fig 4C showing the estimates of the slope of both conditions with and without masking vibrations present. Consistently with Exp. 1, in conditions involving masking vibrations, participants were less precise in discriminating speed and so had a decreased slope. This is also showed in Table 2 where we reported the estimated values of JND in the two conditions (bootstrap estimates and 95% CI).

**Table 2:** Experiment 2: Values of Just Noticeable Difference (JND) in the two experimental conditions. Estimates and 95% CI estimated with the bootstrap methods from the GLMM.

| Estimate | Inferior | Superior | Par | Vibration | Surface |
| --- | --- | --- | --- | --- | --- |
| 0.70 | 0.55 | 0.82 | JND | no | Glass |
| 1.32 | 1.06 | 1.61 | JND | yes | Glass |

#### Contact Force

By means of the force sensor, we measured the force profile for each trial and in each participant. Fig. 1C shows the force profile of one trial in a representative participant. The two peaks of force represent the maximum force in the reference and comparison stimulus. Between the two stimuli, the force is zero because the participant was required to release contact from the plate.

For each participant and trial, we saved the peak of force profile for further analysis. Fig. 5 illustrates the distribution of force peak in the six participants. The median value of force peak ranged from 2.6 to 4.1 N across participants (colored line in the figure). These values are within the range reported in previous studies investigating simulated and natural manipulations [5, 32, 10].

**Figure 5:**
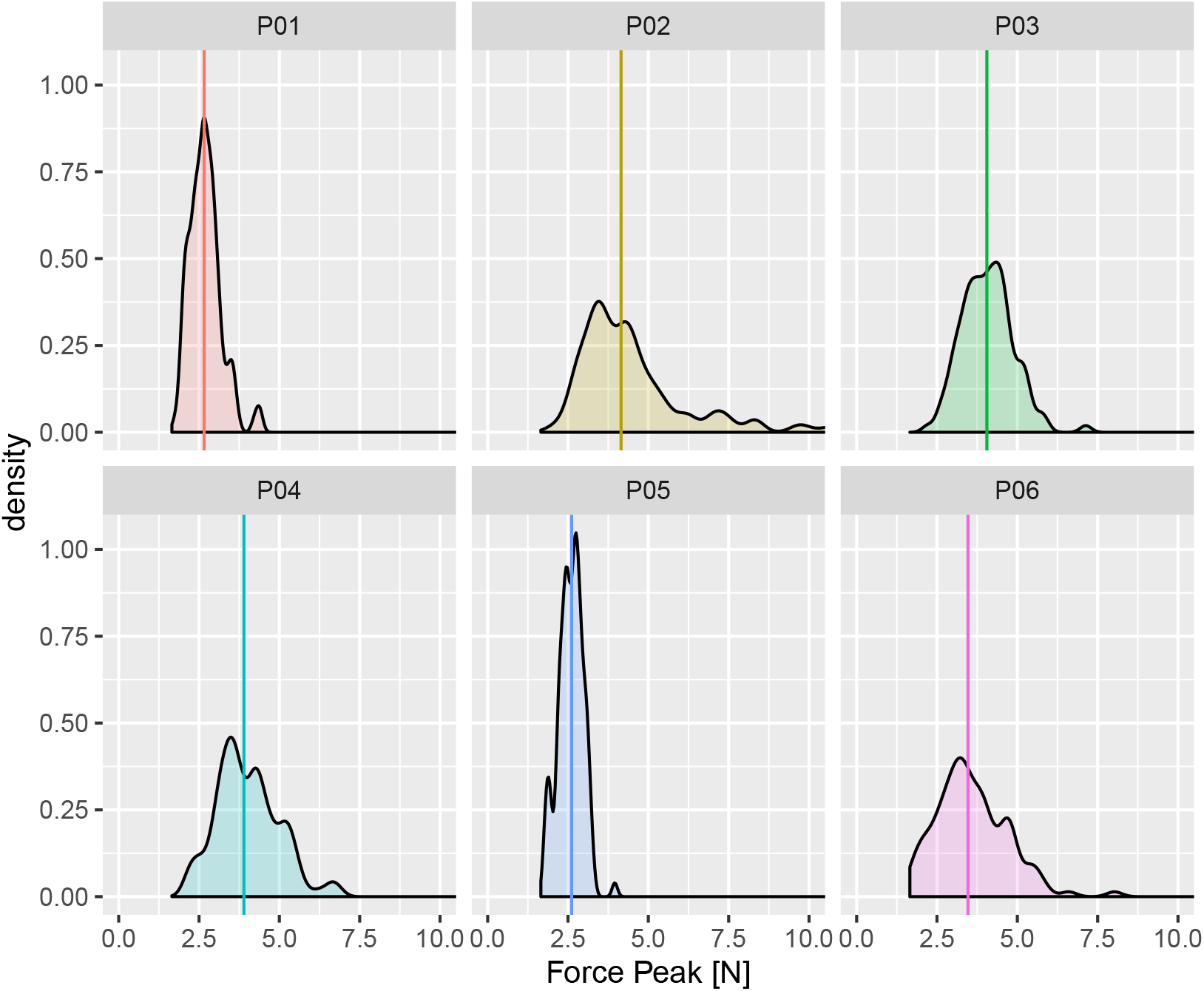
Distribution of the force peak across participants. Values from different participants are illustrated each with a different color. The solid colored lines are the median values.

## Discussion

We investigated the ability of human participants to discriminate the motion speed of a smooth glass plate using touch. Participants were surprisingly good at this: Their responses were only 13% less precise when compared to a control condition, where the motion stimuli were delivered with a plastic plate having a fine texture (Exp. 1). This has important implication for models of motion encoding in touch, because it shows that rich perceptual information is provided even in absence of fine and coarse texture elements. Contact force as measured in Exp. 2 was within a range typical of grasp and manipulation tasks.

The current findings raise the question of which are the tactile cues that convey motion information in absence of detectable texture elements. A recent study suggested that the brain uses a deformation-based mechanism for slip detection on smooth surfaces [10]. Fingertip shearing caused by a stroking glass plate is related to surface displacement, as long as the finger is not fully slipping against the plate. Additionally, the skin in contact with the plate undergoes deformations due to partial slips. In [10], the authors hypothesized that shearing and partial slip may play a role in slip detection [10]. Likewise, speed discrimination on smooth surfaces may be partially based on deformation-rate cues, such as fingertip shearing rate and the onset of the stick-to-slip transition.

Frictional vibrations provide a second class of motion cues in touch. These play an important role in motion detection and for the discrimination of stimulus features like speed, roughness, and texture [6, 22, 15, 17]. Recent findings showed that the responses of Pacinian fibers are strongly modulated by the speed of the moving surface, supporting the hypothesis that the vibration spectrum may provide an auxiliary cue to motion speed [18]. Additionally, during tool use they allow us to discriminate the texture of a surface sensed with a probe, and the position on the probe where it impacts with an object [33, 34]. In a previous study we described a masking effect of vibrations by using either a fine (sandblasted billiard ball) or a coarse texture surface (raised ridges) to deliver the motion stimuli [17]. This masking effect was numerically larger on the fine texture. Likewise, masking electrovibrations impair sharpness perception on touch screen applications [35]. In the current study, we extended these results to speed discrimination on a smooth glass plate. Motion cues based on frictional vibrations and shear force are heuristics, because are not based on *ẋ_i_*, where *x_i_* is the position on the skin of a traceable surface element (e.g., a raised dot). We propose the term *frictional motion cues* to identify these heuristic cues in touch.

Perceptual impairment caused by masking vibrations may be explained either at a mechanical or at a neural level. The small slips between the surface and the skin revealed as skin vibration may provide auxiliary information on surface speed, and these would be masked by the high-frequency vibrations generated by the vibromotor. Masking vibrations may decrease the signal-to-noise ratio of fast adapting fibers, leading to a noisier perceptual response. At a central level, tactile motion and vibratory stimuli activates human middle temporal complex (hMT+) [36, 37]. In particular, vibratory stimuli activate the medial superior temporal area (MST), but not area MT itself [38].

The neural representation of motion in touch could be based on a within-fiber, intensity code [7, 8] and on a between-fiber spatiotemporal code [20]. The latter is defined as the sequential activation of neighboring afferent fibers in response to a traceable feature of the surface, moving across their receptive field [6, 20]. An example of spatiotemporal code is the sequential activation of fast adapting fibers (FA)-I following the motion of a surface with a single raised dot [6]. In the model proposed in [20], tactile motion arises from the responses of the fibers to a moving pattern of skin indentation, based on a spatiotemporal code. This model does not incorporate lateral sliding and the concomitant shear forces. Scanning is mimicked by moving an “image” of the indentation pattern across the skin. The brain may use an intensity code to represent surface motion from the high-frequency vibrations and the gross deformation of the skin [6, 17]. In accordance with the hypothesis of the intensity code, a brush moving across the skin at different velocities produces a modulation in mean firing rate within the single afferent fiber [7, 8]. An intensity code could also account for motion encoding on a smooth surface. Instead, in this case a spatiotemporal code is unlikely for the lack of traceable elements.

These models postulating different motion cues and neural codes are not mutually exclusive. Alike in vision, where an observer is able to perceive the depth of a painting in absence of binocular cues, tactile system can encode and process surface motion also when the stimulus lacks one of the redundant motion cues. For instance, a moving pattern generated by a wave of vibrating pins provide a vivid sensation of a moving surface, even in absence of a net shear force [29, 39]. On the other hand, the results of the current study suggest that vibration and deformation cues may provide rich information to discriminate surface speed.

## Conclusions

This and previous results showed that touch is more than an “imager”, because it is possible to discriminate motion features, such as direction and speed, even in absence of a texture. The different cues—spatiotemporal cues, vibration, and deformation—provide redundant motion information; therefore, they could be integrated for an optimal estimated of tactile motion.

## Author Contribution

**Alessandro Moscatelli:** Conceptualization, Methodology, Visualization, Software, Formal analysis, Writing - Original Draft, Writing - Reviewing and Editing, Supervision. **Colleen P. Ryan:** Conceptualization, Investigation, Data curation, Writing - Original draft preparation, Writing - Review and Editing. **Simone Ciotti:** Software, Methodology, Validation, Visualization, Data curation. **Lucia Cosentino:** Software, Validation, Data curation. **Marc O. Ernst:** Conceptualization, Writing - Reviewing and Editing. **Francesco Lacquaniti:** Conceptualization, Supervision, WritingReviewing and Editing.

## Acknowledgements

This work was supported by the Italian Ministry of Health (Ricerca corrente, IRCCS Fondazione Santa Lucia), Italian Space Agency (grants “‘MARS-PRE” I/006/06/0 and 2019-11-U.0), and Italian University Ministry (PRIN grant “TIGHT”, grant number 2017SB48FP and PRIN grant “NeuAge”, grant number 2017CBF8NJ005). We thank Teresa Ramundo for helping us with data collection.

## Footnotes

1 The estimated values of *η*_5_ is slightly smaller than the difference between *β*_1(*glass*)_ and *β*_1(*plastic*)_. This is due to the choice of the random-effect parameters in Eq. (2).

